# Backyard proteomics: A case study with the black widow spider

**DOI:** 10.1101/2025.04.11.648442

**Authors:** Tarsh Shah, Jackson A. Fitzpatrick, Benjamin C. Orsburn

## Abstract

Nearly all methods of mass spectrometry-based proteomics rely on knowing the proteome of the species. In less studied organisms without annotated genomes, it can seem impossible to perform proteomic analysis. In this study we sought to answer the question - does enough information exist to do proteomics on any organism we want? As a case study we started with material available due to an infestation of a home with black widow spiders. Thanks to the recent publication of an annotated genome for one species of black widow spider, we were able to identify 5502 protein groups and assign putative annotations using ortholog mapping. We also demonstrate that had we not had this resource, over 2000 proteins could be identified using other available spider genome annotations, despite their unrelatedness. Moreover, regardless of spider proteome used, proteins annotated as toxins were almost exclusively observed in the main body of the mature female black widow spider. Overall, these results provide a draft proteome map for the black widow spider and valuable data for validating machine learning models, while also suggesting that the door to insightful quantitative proteomics may already be open for millions of less studied organisms. All raw and processed proteomic data are available through the ProteomeXchange repository as accession PXD051601.

**TOC Graphic:** 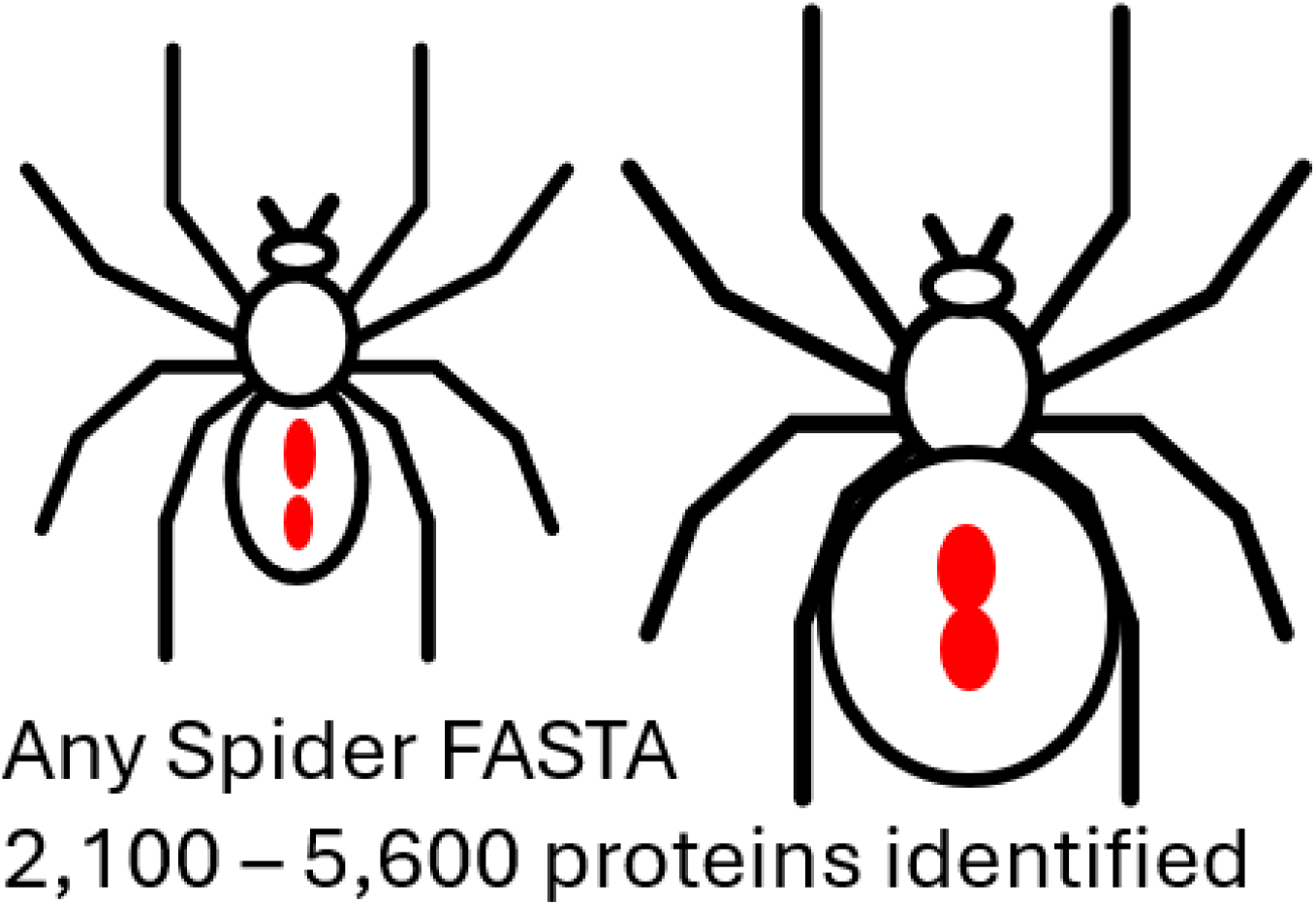

## Introduction

Today, quantitative proteomics can be performed by a dizzying number of technologies, many of which can employ multiple techniques. Liquid chromatography mass spectrometry (LCMS) can acquire untargeted data with either data dependent acquisition, relying on on-board computers for on-the-fly target acquisition, or data INdependent acquisition where you fragment it all and let the algorithms sort it out.^1^ You can also use LCMS systems to specifically target what you’re interested in or mix and match these methods for various semi-targeted methods.^2,3^ Older and newer techniques exist where proteins are targeted and measured by antibodies, through protein arrays or by oligonucleotide amplification.^4^ You can even estimate protein presence and concentration within a narrow dynamic range through aptamer binding assays.^5^ While a tipping point may be approaching soon for so-called “next generation” techniques thanks largely to their impressive throughput capabilities^6^, most proteomics labs in the world probably rely on LCMS technologies at the date of this writing. While the earliest techniques in LCMS based proteomics relied heavily on *de novo* sequencing tools, the productivity of the field increased exponentially in the wake of the resources produced by genomic sequencing technologies.^7^ Today, nearly every LCMS proteomic software tool relies on curated libraries of protein or peptide sequences derived from annotated high quality genome assemblies. The form of this input is generally a fasta file, which is a flat format of protein sequences, or spectral libraries, which are experimentally obtained spectra matched to the same fasta.^2^ The productivity of LCMS proteomics again increased substantially due to the application of deep learning tools which can now predict peptide fragmentation patterns from the curated knowledge of the past.^8,9^ While these deep learning tools are often described as “library-free” this is somewhat of a misnomer as protein sequence input is still required from which peptide masses and fragmentation patterns are modeled and predicted.

The end result of these advancements is that LCMS-based proteomics has near universal methods that can provide proteomic data on practically any sample type, but this is largely limited to organisms with annotated genome assemblies. At the end of 2024, NCBI had annotated 1286 eukaryotic species^10^, though this does not mean all 1286 have had proteomic analysis. An estimated 8.7 million distinct species of eukaryotic life exist on earth today, meaning these annotated eukaryotic genomes provide minimal coverage of the tree of life.^10^ Some groups, like mammals, have more annotated genomes relative to the number of species, but even then some clades have more annotated genomes than others.^11^ This paucity of unannotated species genomes is mirrored in published proteomic data. As of February of 2025, humans comprise 47 % of the roughly 40 000 mass spectrometry proteomic datasets on ProteomeXchange, followed by mouse at 17 %, then a steep drop off to 2.9 % as the other classic model organisms comprise the top 10 species.^12^ We will not diminish the importance of human research, but emphasize the diversity of species in the world around us that lack proteomic analysis and that this presents an opportunity for researchers everywhere.

There are different reasons to perform proteomic analysis on a given species, which can include everything from developing drugs against that organism^13^ through learning how to synthesize molecules it produces^14^, but it also could simply be curiosity of the unknown. But should you forgo exploratory analysis when there is not an annotated genome assembly? Though the specific annotated genomic landscape (i.e., what is the closest related species that is annotated) of a species will be unique, we suggest that proteomic analysis should be considered viable, and that the data will be valuable in the future as more genomes are sequenced and annotated. For this example, we performed proteomic analysis on a black widow spider (*Latrodectus* spp). There are around 52 000 spider species^15^ and only 46 Araneae have reference genomes on NCBI, of these, only 14 have annotated genome assemblies (as of August 2024). The black widow spider is in the Theridiidae family which has one annotated genome assembly on NCBI, *Parasteatoda tepidariorum*. While multiple proteomic studies have focused specifically on the toxins produced by black widow spiders,^16,17^ no comprehensive proteomic analysis has been performed on the whole spider. Based on the availability of specimens within persistent infestation of a home in Pennsylvania, we have sampled two individuals: a single intact spiderling and a mature spider, with the latter in a partially homogenized state due to the utilization of an aluminum baseball bat by a nervous mass spectrometrist during sample collection. In total, six separate anatomical regions could be excised from these organisms for proteomic analysis. After analysis, the Western black widow spider (*Latrodectus hesperus*) genome assembly was published^18^ and upon request authors provided the fasta for the annotated genome, albeit without useful protein names. Using this *L. hesperus* annotated genome, we provide here a draft proteome map of the black widow spider but also compare to proteomic analysis if we only had the 14 other spider genome annotations. The analysis using the *L. hesperus* annotated genome was unsurprisingly superior, but proteomic analysis using the 14 NCBI spider genome annotations provided valuable results, including putative biological insights. These results highlight that proteomic analysis of specimens from the world around us (even our backyard) can be useful even when a species-specific fasta does not exist.

## Methods

### Spider collection

Following multiple attempts to collect protein from spiders homogenized by aluminum baseball bat (Marucci F5, Baton Rouge, Louisiana), an author missed striking female spider directly. This specimen was only partially homogenized and likely incapable of murdering the author who made undignified sounds while transferring her into a 15 mL centrifugation tube (Falcon). A second immature spiderling was captured fully intact using an eco-friendly bug vacuum for kids (Nature Bound Toys, Denver, Colorado). The spiderling was transferred to a 15 mL centrifuge tube and both samples were stored at -20 °C before transporting to the lab on wet ice for long term -80 °C storage.

### Spider protein sample preparation

The *Latrodectus* specimens were aseptically transferred into a 15 mL falcon tube for further processing. Upon thawing, the specimens were dissected by best effort into discernible sections. For the intact spiderling, four distinct regions were excised. For the adult spider for which capture was more challenging and resulted in partial homogenization, only two regions could be confidently collected and were designated as “body” and “legs”. Homogenization was carried out using a bead mill (BeadBug 3, Benchmark Scientific, Sayreville, New Jersey) in 2 mL bead chambers, employing 30 s pulses of creepy spider parts in 700 μL of 1x S-Trap lysis buffer (5 % sodium dodecyl sulfate; Protifi, LLC). Following homogenization, the resultant lysates were centrifuged at 13 000 x *g* for 3 min to facilitate clarification, thereby separating the liquid fraction protein fraction from solid and creepy black chunks of chitin.

The clarified homogenates were prepared by suspension trapping^19^ using S-Trap Mini spin columns (Protifi) with minor modifications of established manufacturer protocols. Protein samples solubilized in 1x S-Trap lysis buffer were reduced with dithiothreitol (DTT; Thermo Fisher; final concentration: 10 mmol/L) at 60 °C for 20 minutes followed by alkylation with iodoacetamide (IAA; Thermo Fisher; final concentration: 40 mmol/L) at room temperature in a dark drawer for 20 min. A total of 40 μL of clarified homogenate was acidified with 12 % phosphoric acid and diluted 1:10 (volume fraction) in S-Trap binding buffer (80 % LCMS grade methanol with 100 mmol/L triethylammonium bicarbonate, pH 8.0) and centrifuged at 2000 x g for 2 min to bind the proteins to the S-Trap column. The bound proteins were then washed three times with 300 μL of binding buffer. Finally, the clean bound proteins were digested by the addition of 10 μg of MS-grade trypsin (Thermo Scientific) per sample in a total of 80 μL of 100 mmol/L triethylammonium bicarbonate solution, resulting in the enzymatic cleavage of proteins into peptides. Following digestion overnight at room temperature, peptides were eluted with 0.2 % formic acid (volume fraction) in water and dried under vacuum centrifugation (Eppendorf). Subsequently, they were resuspended in 200 μL of 0.1 % formic acid (volume fraction) in water, and their concentrations were quantified using the Pierce Quantitative Colorimetric Peptide Assay kit (Thermo Scientific) following vendor protocol.

### LCMS Analysis

The peptides were diluted and 400 ng peptides per sample loaded onto each individual EvoTip (EvoSep, Odense, Denmark) and injected into timsTOF FleX mass spectrometer (Bruker) through Evosep One (Evosep, Odense, Denmark) liquid chromatography system. The 30 sample per day (SPD) separation program was used which is a 44 minute separation on a 15 cm x 150 μm C-18 column with 1.5 μm particle size (PepSep, Odense, Denmark). The outlet column was connected through a “zero dead volume” union to a CaptiveSpray emitter with a 10 μm emitter tip. Eluted peptides were analyzed with diaPASEF using a method constructed from the default “diaPASEF short gradient” in Bruker TIMSControl 4.0.

### Data Analysis

The resulting mass spectrometry (.d) data files were automatically converted by using a batch command in Bruker Hystar to HTRMs with the appropriately named “HTRMS Converter” software (Biognosys). The files were then transferred to a central server for processing in SpectroNaut 19. Search settings for SpectroNaut followed the default DIA+ “deep” workflow for TIMSTOF data. Briefly, SpectroNaut determined the appropriate precursor and fragment ion tolerances with carbamidomethylation of cysteines set as a static modification. Oxidation of methionine and acetylation of the protein N-terminus were the sole dynamic modifications considered, with a maximum of 3 dynamic modifications considered on each peptide sequence. Up to 2 missed tryptic cleavage sites were considered. For downsteam analysis a pivoted report for the proteins and peptides were separately exported using the default templates with the additions of the protein description and gene identifiers, which are strangely absent in both default templates. A protein fasta of the recently published genome assembly of the Western Black widow spider (*Latrodectus hesperus hesperus*; taxonomy ID: 256737) was obtained from the authors.^18^ This genome assembly is available on Genbank as ASM3797512v2 (GCA_037975125.2) but has not been annotated by RefSeq, which is why we used the author supplied fasta. This fasta was used for *L. hesperus* specific searching is labeled 68345-CDS-prot.fasta (34 991 sequences). The original CDS matches and linked annotations are provided as **Supporting Information 1** while the other 14 spider fasta are provided **Supporting Information 2**. Note that the *L. hesperus* fasta was provided mis-labeled within fasta headers as *Hesperus amabilis*, but this was simply a mistake in the system.

## Data Availability

Peptide level analyses and all .fasta files exceed the maximum allowable file size for this journal and have been permanently published on FigShare and are available at the following https://doi.org/10.6084/m9.figshare.28462295.v1. All raw and processed proteomic data are available through the ProteomeXchange repository as accession PXD051601.

### Ortholog mapping of protein identifications

This fasta used had no useful information for protein descriptions in the fasta header, and so ortholog mapping was used to assign putative identifications to allow proteomic results to be interpreted. Briefly, make_subset_DB_from_list_3.py and db_to_db_blaster.py from PAW_BLAST GitHub repo were used locally to BLAST (v.2.11.0+) each of the identified protein entries against a fasta comprised of all the UniProt entries under the taxonomy Araneae (Spider Order; taxonomy ID: 6893) from the 2024_07 UniProt release (614 548 entries). This UniProt fasta was used because the naming and descriptions was more complete than using any of the 14 NCBI annotated genome assemblies. We expect there to be an official NCBI annotated black widow genome assembly in the future that will have curated protein names and descriptions, adhering to RefSeq levels of quality. These approximately 5500 mapped protein names were not checked for poor or no homology, but instead were used for descriptive purposes and should be considered putative.

### Analysis of toxin proteins

All output protein reports were exported as .tsv files from SpectroNaut using the default protein pivot report option with the addition of the Protein Group Description column enabled. The .tsv files were combined into a single Excel file with 16 sheets and each sheet was filtered in place on the term “toxin”. The resulting lists were combined into a single Excel sheet and the RAC-1 proteins were manually deleted as this ubiquitous Eukaryotic protein description contains the letters “toxin” but is clearly unrelated. The sheet was then copied into the Broad Morpheus visualization program using default parameters for heatmap creation (https://software.broadinstitute.org/morpheus).

## Results and Discussion

### Generating a draft proteome map

The majority of spider research, including specifically the black widows, is focused on silk ^17,20^ and toxins^21,22^, largely ignoring the rest of the spider. As of February 2025, ProteomeXchange lists 29 data sets of spider proteomes, with only 2 including multiple tissues, with the obvious exception of the data published with this study. Though this study was advantageous in nature, it still offered an opportunity to provide a draft proteome for the black widow spider due to multiple tissue samples. Though unavailable until well after initial data analysis, the *L. hesperus* annotated genome allowed for peptide identifications in the DIA mass spectrometry data. The identifications in each tissue ranged from 2,422 to 39,280 peptides, with the most identified in the adult body and the least in the small head of the immature spiderling. These in turn resulted in 5,608 protein identifications across the experiment, ranging from 1135 in the spiderling head to 5602 in the legs of the adult spider. Due to sample harvesting samples with the adult spider previously mentioned, the two sections contained a homogenate of more anatomical regions than others. It should also be noted that the spiderling was quite small and excision of the head was difficult and resulted in limited peptides for analysis. The gene annotation itself in the *L. hesperus* annotated genome seemed accurate in the sense that it explained the collected data well enough, but the gene identifications themselves were not informative. To overcome this limitation we mapped each identified protein to its Araneae (Spider Order; taxonomy ID: 6893) ortholog in order to provide putative identifications. This complete proteome map with putative protein IDs and relative abundance across all tissues is available in **Supporting Information 1**. The dynamic range of each tissues proteome is plotted and the top 5 most abundance proteins are available in **Supporting Information File 4: Figures S1 to S6**, and the underlying data was derived from **Supporting Information File 1: Table S1**. Finally, we have broken up the identifications by tissue (and by individual) and sorted from highest to least abundance. This can be interesting to see what the most abundant proteins are in each tissue, but we acknowledge that without more replicates (both biological and experimental) that any subtle differences in abundance should be interpreted with caution. Overall, to our knowledge this is the first multi-tissue proteomic analysis of this species.

### Comparison to non-species specific search results

A major roadblock to performing proteomics in many species is the lack of a species-specific annotated genome to search the mass spectrometry data against. During this study we were able to use a newly annotated genome, but we wanted to examine how the results might have looked without a species-specific annotated genome. To demonstrate this we analyzed the data using each of the 14 spider genome annotations (**Table 1, Supporting Information 2**). Unsurprisingly, the results of using the species-specific genome annotation was better than using these other species with 50 699 experiment-wide peptide identifications. Using identified peptides as a comparison measure we found the worst performing spider genome annotation to be *T. clavipes* with 8274 IDs, 16.3 % of peptide IDs versus the species-specific genome, and the best performing spider to be *P. tepidariorum* (the common house spider) with 12 361 peptide IDs, 24.4 % on peptide IDs versus the species-specific genome. Interestingly, in the case of the *P. tepidariorum* results, of the 12 361 peptide identifications, 10 151 peptides were shared with the species-specific search. These differences seem to recapitulate the relatedness of these species (both the black widow and common house spider are in the Theridiidae family), which re-iterates that when using a non-species specific genome annotation, it is best to use the closest relative possible.

**Table 1.** A summary table of the number of peptide groups, proteins and protein groups obtained when files were searched against each .fasta separately.

| Fasta file used | Taxonomy ID # | Peptides | Proteins | Protein groups |
| --- | --- | --- | --- | --- |
| *Araneus ventricosus* | 182803 | 9913 | 3829 | 2464 |
| *Argiope bruennichi* | 94029 | 10752 | 3442 | 1956 |
| *Trichonephila clavata* | 2740835 | 10510 | 2889 | 2344 |
| *Trichonephila clavipes* | 2585209 | 8335 | 2542 | 1968 |
| *Trichonephila inaurata madagascariensis* | 2747483 | 9657 | 2377 | 1980 |
| *Nephila pilipes* | 299642 | 10471 | 2754 | 2156 |
| *Parasteatoda tepidariorum* | 114398 | 12361 | 4396 | 2347 |
| *Uloborus diversus* | 327109 | 9962 | 2777 | 2028 |
| *Stegodyphus dumicola* | 202533 | 10003 | 3083 | 1974 |
| *Stegodyphus mimosarum* | 407821 | 9805 | 2098 | 2044 |
| *Oedothorax gibbosus* | 931172 | 9611 | 2610 | 2045 |
| *Caerostris darwini* | 1538125 | 10658 | 2542 | 2110 |
| *Caerostris extrusa* | 172846 | 10272 | 2143 | 1902 |
| *Larinioides sclopetarius* | 280406 | 10897 | 3786 | 2250 |
| ***Latrodectus hesperus* (CDS)** | 256737 | 50,699 | 5608 | 5502 |

### High relative concentrations of toxin proteins are observed in the mature female spider

Finally, we performed an exploratory analysis using the species-specific results for toxins present across tissues. In every case, each annotated fasta file used provided a minimum of 7 protein group annotations with a description containing the world “toxin”. Following removal of RAC1 protein, which are highly conserved RAS-related botulinum toxin substrate proteins and unrelated to spider toxins, a minimum of six toxin were identified in all analyses. (**Supporting Information 3**). When visualized the abundance of these proteins using row normalized heatmaps (**Figure 2**) we find that nearly all were primarily observed in the mature female spider body, with lower relative levels in the female legs. Toxin proteins were only sporadically detected in the immature spiderling. Recent focused work on Latrodectus toxins described the bite of a mature spider as an “arsenal” of various toxins with specificity to a broad range of organisms. Some of these proteins appear to only affect mammals while others have broad impacts across all vertebrates. Immature spiderlings do possess toxins, and extracts from immature spiderlings have demonstrated toxicity in some animal testing. It is thought, however, that while spiderlings have toxins for insects for paralyzing and consuming them, the full arsenal of toxins are reserved for mature females when protecting their eggs and young.^23^ The proteomic data appears to support this hypothesis, regardless of the FASTA database utilized in the analysis of these files.

**Figure 1.**
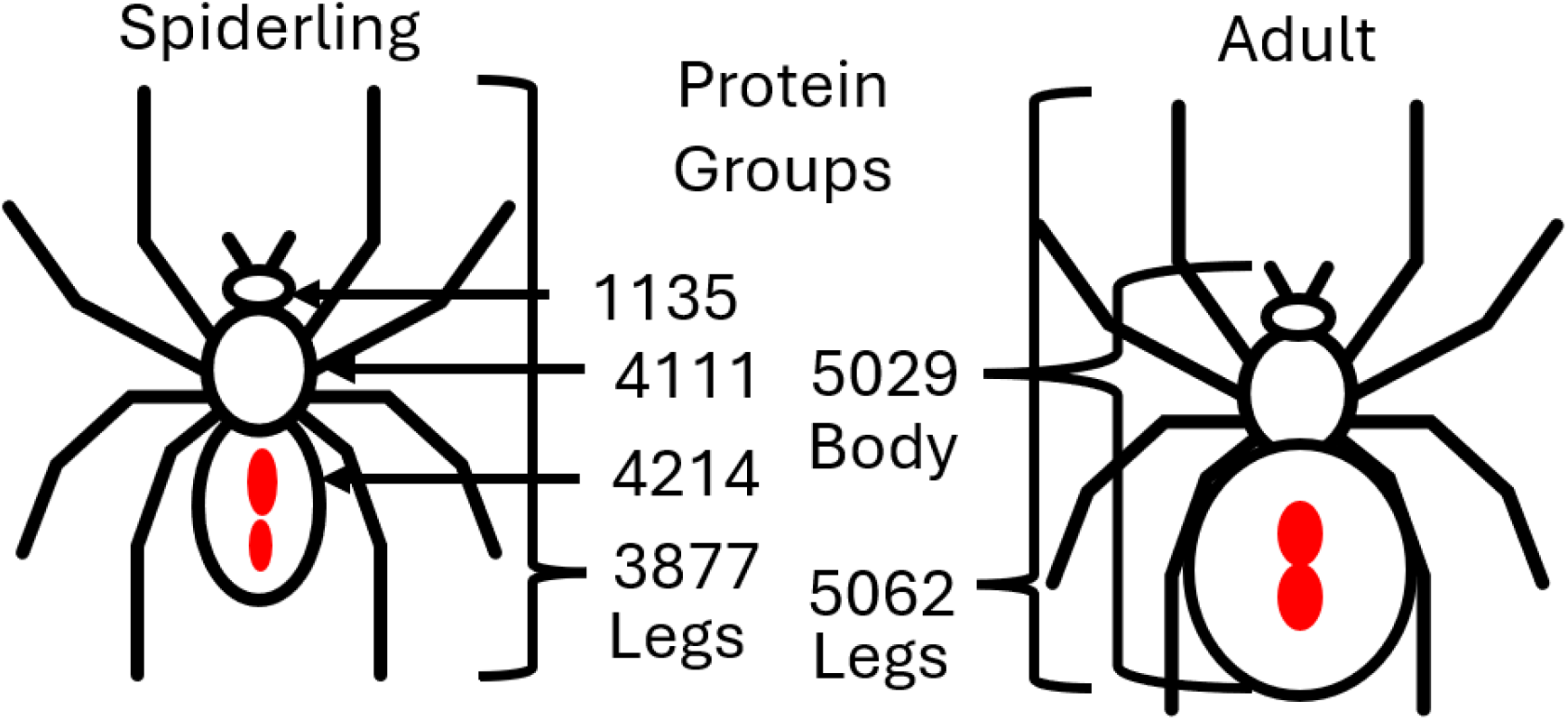
A schematic representing the samples which were taken for proteomics and the number of protein groups identified in each sample when using translated coding DNA sequences from the Western Black Widow genome.

**Figure 2.**
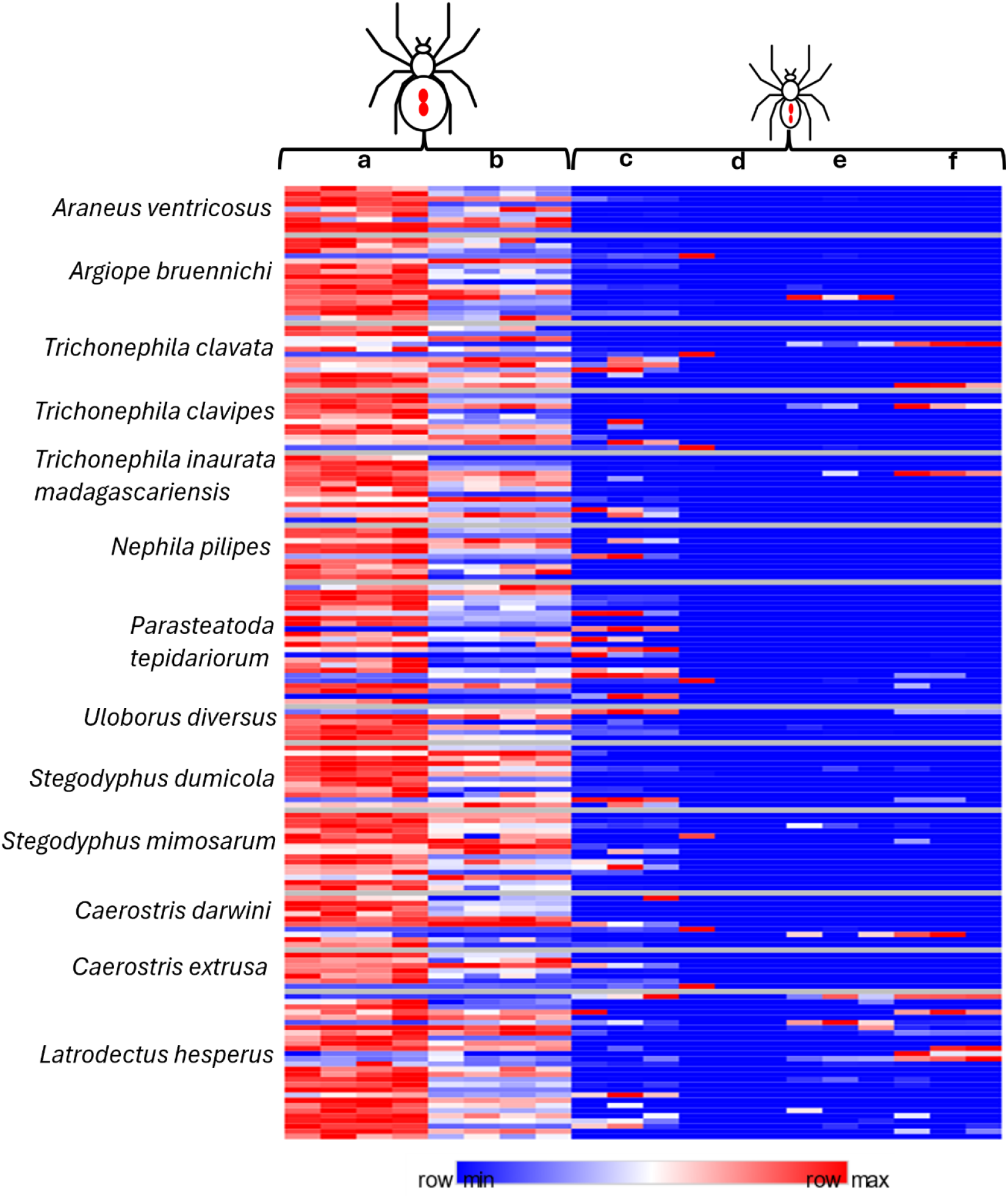
A heatmap demonstrating the relative abundance of proteins annotated as toxins in each FASTA database for each spider region analyzed. Mature female body (a) and legs (b). Spiderling abdomen (c), head (d), legs (e) and sternum (f).

Though these results are exploratory and are more for demonstration, it does emphasize that even without the species-specific spider genome we would have been able to make putative biological conclusions. In either case these results would require additional work to confirm, but undoubtedly they are hypothesis generating.

## Limitations

The 14 non-black widow spider annotated genomes are a combination of four official RefSeq annotations and 10 user submitted. This can create a difference in quality that makes any species comparisons difficult.

Also, because we did not go through the ortholog mapping of over 5000 black widow proteins, there are undoubtedly duplicate protein hits or missing identifications since the fasta used likely was not only canonical/non-redundant gene products and contained putative assignments and fragmented sequences. We expect that in the future when RefSeq releases the black widow genome annotation using this same genome assembly, that the protein identifications will be more useful and this search and compilation can be re-performed. Still, this does not negate the findings and comparisons here, especially as our goal was not to drive new biological insight but instead to provide a draft resource to be improved upon in the future.

## Conclusions

There is no question that this is an exciting time to be doing proteomics.^24^ The field may even be emerging from under the shadows of genomics and receiving mainstream attention thanks to recent success stories in the clinic and new datasets featuring thousands or tens of thousands of patient proteomes.^6^ New reagents and smart methods even allow the proteomic analysis of body fluids with extremely high dynamic ranges in protein concentration which have traditionally been the hardest things to study with proteomics.^4^ It seems no coincidence that most of the big stories have been in human proteomics, which is supported by the availability of both small curated genomes and vast libraries of human mutations.^25^ Recently we have even seen new resources to address population level genetic variations to enable comprehensive studies of people with racial and ethnic backgrounds under-represented in current databases.^26^ When you step outside of human studies, however, many of these resources have no parallel and seem to suggest that you simply can’t do proteomics outside of a few model organisms. However, it should be noted that thousands of fasta databases are available for analysis today and that they can be readily created with a little effort from the genomic data that has been acquired or is being output on sequencers today.

Even in the absence of species-specific fasta databases, there are many reasons to just go ahead and do the proteomics if and when you can. The first is that there is likely sufficient protein sequence homology in the fasta files that are out there to make matches. In the presented case, every spider fasta allowed the identification of over 2,000 proteins, and every library correctly identified the body of the mature female spider as the one with the highest relative concentration of toxin proteins. With freely available tools for protein annotation we were able to match over 5500 predicted protein coding regions to spectra and assign putative annotations to over 90 % of these protein level matches using orthology mapping to all available Araneidae sequences on UniProtKB. The publication of the black widow genome assembly occurred by coincidence within the same time frame as our study^18^ and the author’s annotation was graciously provided by the authors. If this species-specific genome annotation hadn’t been published before we completed our analysis, we simply could have waited. Someone would have inevitably sequenced the genome of a black widow spider. If we wait a bit longer it is likely that a fully annotated reference fasta for this important and creepy spider will become available for an even more complete analysis of these publicly available proteomics files we have generated. However, while we identify far fewer proteins using non-species specific databases, we find that those results can still be biological meaningful and did not require waiting for perfection. If there is a take away from this backyard proteomics study, it is that the correct time to do proteomics is whenever you have protein to analyze. Though it may seem frivolous to engage scientific curiosity in the world around you, there is a legitimate need for data from a broader diversity of species beyond model organisms. State of the art algorithms of peptide spectra prediction and *de novo* are using public data for training and validation, with nearly half of public data sets on PRIDE being human derived.^12,27^ There is value in generating data from species that aren’t human or even model organisms, including areas of the tree of life that are nearly completely lacking, such as insects and specifically spiders. These data could be used in training, validation, or in benchmarking algorithms.

## Supporting information

The supporting information is available free of charge at:

**Supporting information 1** is an Excel file containing the original CDS protein level matches in sheet 1 as well as the assigned annotations and annotation scoring parameters combined with the protein level report as sheet 2.

**Supporting information 2** is an Excel file containing a summary table as well as individual sheets for all quantitative protein values for all samples against each respective .fasta file.

**Supporting information 3** is an Excel file containing each toxin protein identified when using each respective .fasta file along with the quantitative source data used to generate Figure 2.

**Supporting information 4** is a word document containing **Supplemental Figures S1 – S6**.

## Acknowledgements

We would like to thank Dr. Colten Eberhard and Ahmed Warshanna for training T.S. in proteomics sample preparation and for support in both instrument operation and data analysis. We thank Dr. Trevor Krabbenhoft for sharing their annotated black widow spider genome fasta. We would like to thank Marina Pominova and Wout Bittremieux for their insight in the value of this type of data in machine learning training, validation, and benchmarking. Finally, we would like to thank the support of Ben Neely and the National Institute of Standards-Charleston.

## Funding

This study received no external funding.

## Notes

https://proteomecentral.proteomexchange.org/cgi/GetDataset?ID=PXD051601

### Competing Interest Statement

The authors have declared no competing interest.

