## Supporting Information 4 for "Backyard proteomics: A case study with the black widow spider"

Supplemental Figure S1. Average protein abundance versus rank of proteins identified in the adult body.

Supplemental Figure S2. Average protein abundance versus rank of proteins identified in the adult limbs.

Supplemental Figure S3. Average protein abundance versus rank of proteins identified in the spiderling abdomen.

Supplemental Figure S4. Average protein abundance versus rank of proteins identified in the spiderling head.

Supplemental Figure S5. Average protein abundance versus rank of proteins identified in the spiderling legs.

Supplemental Figure S6. Average protein abundance versus rank of proteins identified in the spiderling sternum.

**Supplemental Figure S1. Average protein abundance versus rank of proteins identified in the adult body.** Ranked top 5 most abundant proteins (starting with most abundant): Fatty acid-binding protein, Myosin heavy chain, muscle, Actin, clone, Uncharacterized protein, Cuticle protein 16.8. Protein identifications were mapped from *L. hesperus* genome annotation-based identifications to available Araneae UniProtKB protein entries as described in methods and available in Supporting Information 1. These identifications are putative and no homology threshold was used.

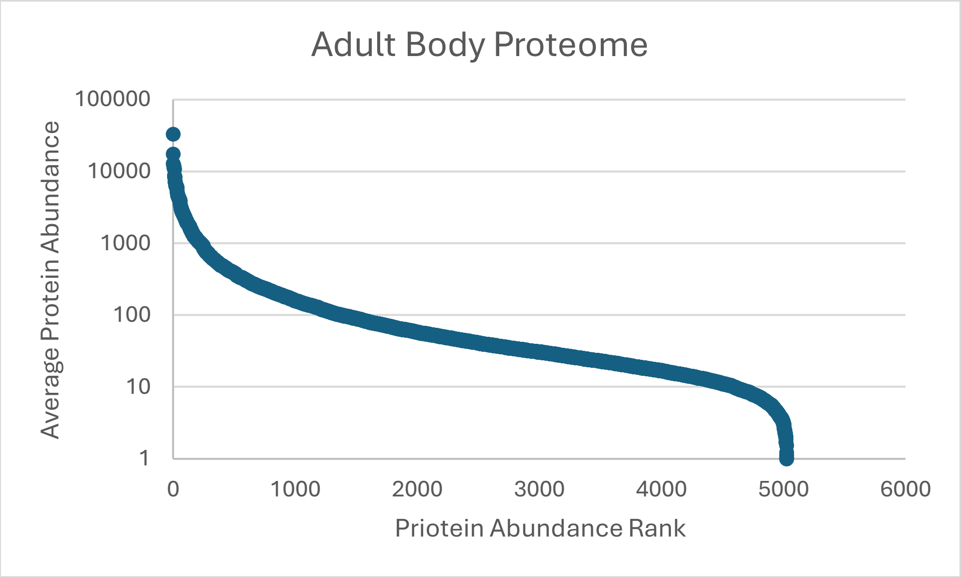

**Supplemental Figure S2. Average protein abundance versus rank of proteins identified in the adult limbs.** Ranked top 5 most abundant proteins (starting with most abundant): Adult-specific rigid cuticular protein 12.6 (Fragment), Glyceraldehyde-3-phosphate dehydrogenase, Papilin, Fructose-bisphosphate aldolase, and Actin, clone 403. Protein identifications were mapped from *L. hesperus* genome annotation-based identifications to available Araneae UniProtKB protein entries as described in methods and available in Supporting Information 1. These identifications are putative and no homology threshold was used.

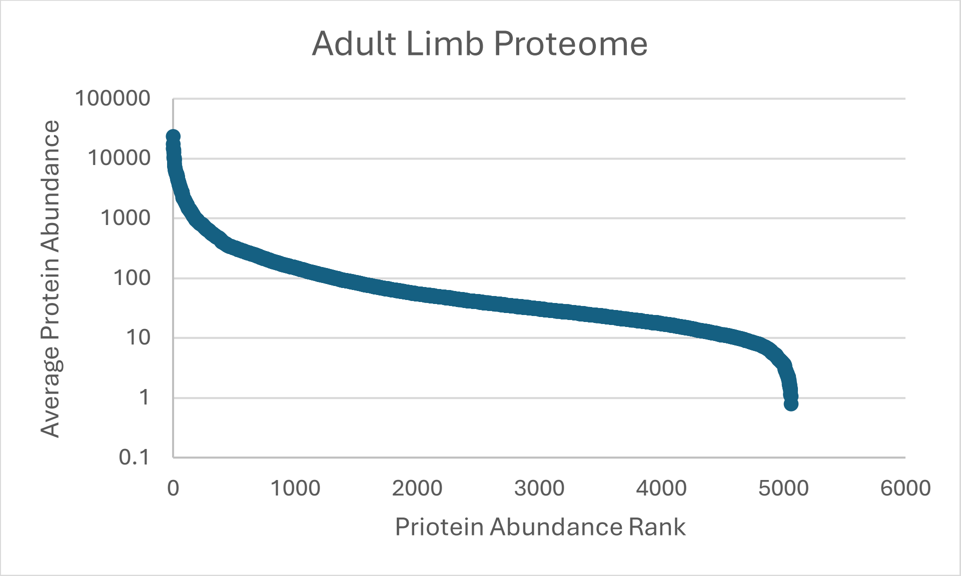

**Supplemental Figure S3. Average protein abundance versus rank of proteins identified in the spiderling abdomen.** Ranked top 5 most abundant proteins (starting with most abundant): Aciniform spidroin 1, Uncharacterized protein S, Cuticle protein 16.8, Polyubiquitin-A, and Rab-GAP TBC domain-containing protein. Protein identifications were mapped from *L. hesperus* genome annotation-based identifications to available Araneae UniProtKB protein entries as described in methods and available in Supporting Information 1. These identifications are putative and no homology threshold was used.

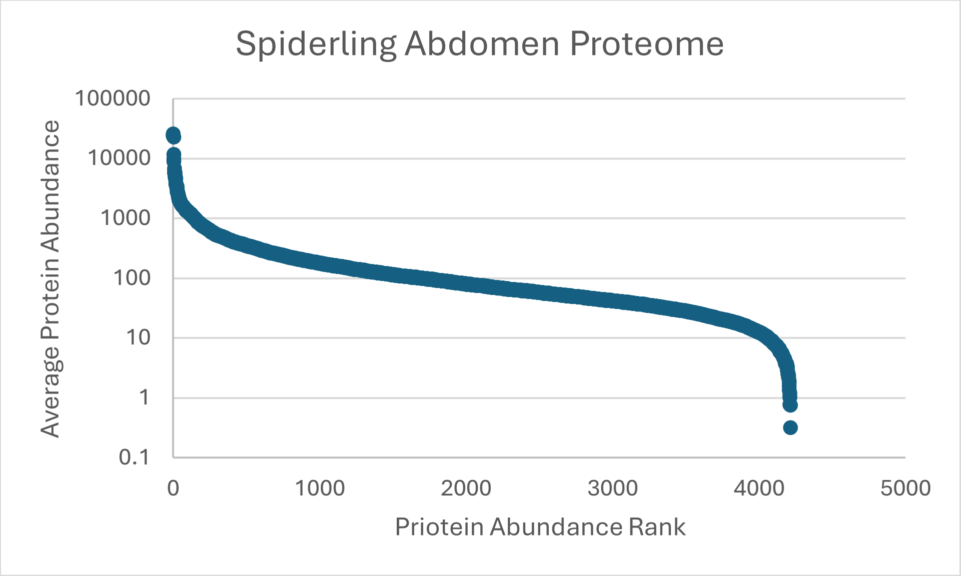

**Supplemental Figure S4. Average protein abundance versus rank of proteins identified in the spiderling head.** Ranked top 5 most abundant proteins (starting with most abundant): ADP/ATP translocase (Fragment), Histone H1E, Putative cytochrome c (Fragment), Uncharacterized protein, and Unconventional myosin-Id. Protein identifications were mapped from *L. hesperus* genome annotation-based identifications to available Araneae UniProtKB protein entries as described in methods and available in Supporting Information 1. These identifications are putative and no homology threshold was used.

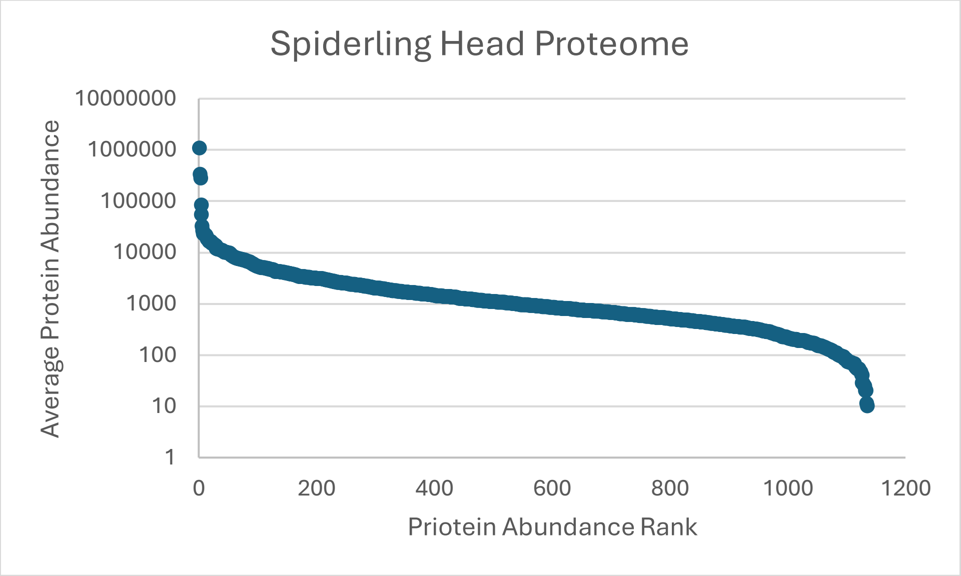

**Supplemental Figure S5. Average protein abundance versus rank of proteins identified in the spiderling legs.** Ranked top 5 most abundant proteins (starting with most abundant): Glyceraldehyde-3-phosphate dehydrogenase, Adult-specific rigid cuticular protein 12.6 (Fragment), Myosin heavy chain, striated muscle, Hemocyanin F chain, and Myosin heavy chain, muscle. Protein identifications were mapped from *L. hesperus* genome annotation-based identifications to available Araneae UniProtKB protein entries as described in methods and available in Supporting Information 1. These identifications are putative and no homology threshold was used.

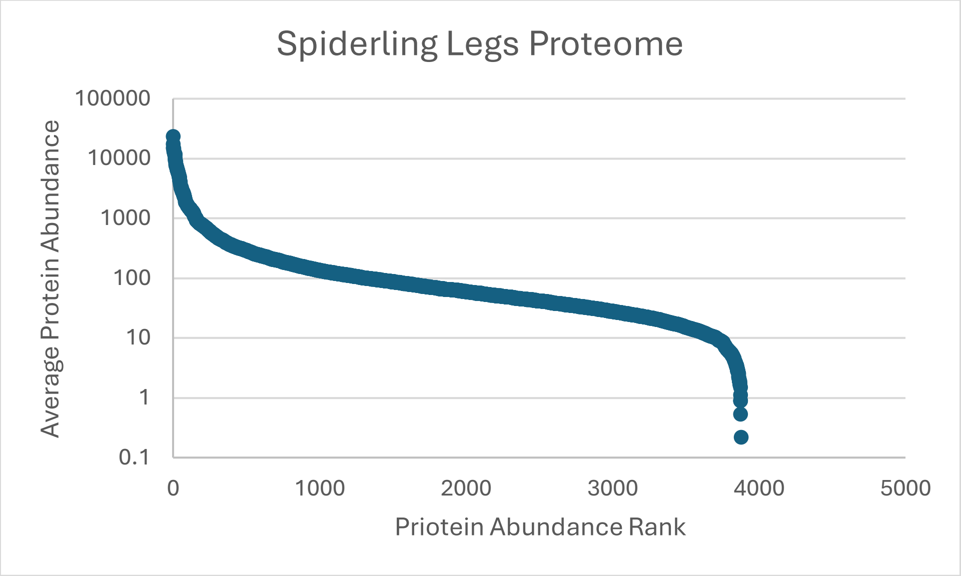

**Supplemental Figure S6. Average protein abundance versus rank of proteins identified in the spiderling sternum.** Ranked top 5 most abundant proteins (starting with most abundant): Caspase like protein, Rab-GAP TBC domain-containing protein, Glyceraldehyde-3-phosphate dehydrogenase, Myosin heavy chain, striated muscle, and Uncharacterized protein. Protein identifications were mapped from *L. hesperus* genome annotation-based identifications to available Araneae UniProtKB protein entries as described in methods and available in Supporting Information 1. These identifications are putative and no homology threshold was used.

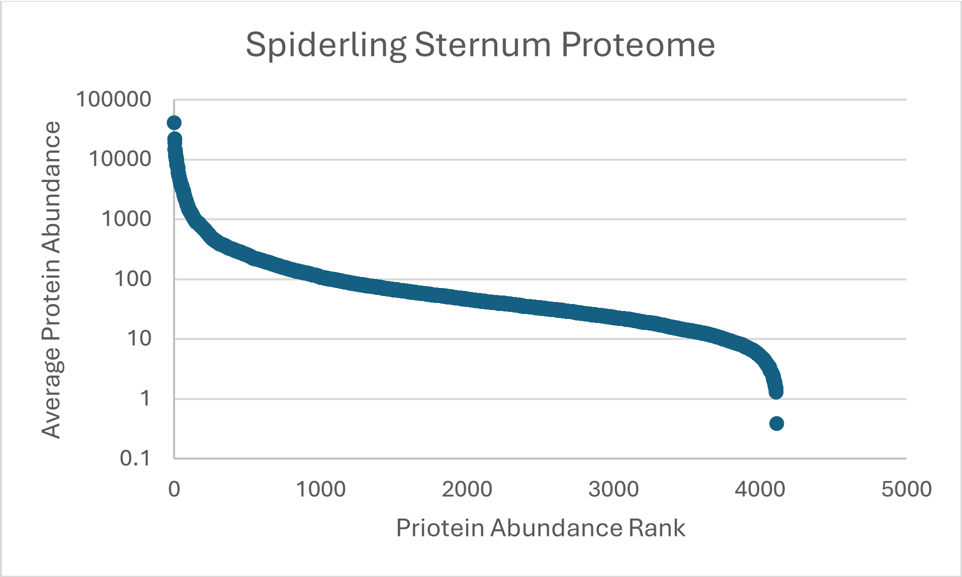
